# Single-Channel EEG Artifact Identification with the Spectral Slope

**DOI:** 10.1101/2023.11.12.566749

**Authors:** Melissa C. M. Fasol, Javier Escudero, Alfredo Gonzalez-Sulser

## Abstract

Electroencephalogram (EEG) signals are a valuable recording technique to diagnose neurological disorders and identify noninvasive biomarkers for clinical application, however, they are vulnerable to various artifacts. It is difficult to define exact parameters which efficiently distinguish artifacts from neural activity, and thus cleaning EEG data often relies on labor-intensive visual scoring methods. While signal processing techniques to remove artifacts exist, many state-of-the-art techniques are designed for multivariate signals, which can be challenging to implement in recording setups with few electrodes. We demonstrate how the spectral slope - a method previously used to distinguish between conscious states by linear regression of the logarithmic EEG power spectra - can also be used to identify epochs contaminated by recording artifacts in rat EEG recordings and propose this as a first pass artifact detection method. We computed the mean spectral slope for both ‘clean’ and ‘noisy’ epochs and compared the distributions among individual recordings to determine whether the decision threshold should be dynamic or fixed. We found no significant difference between the mean of these distributions and determined that a spectral slope threshold of -8 *μV* ^2^*/Hz* was effective at identifying noisy epochs across all recordings. The accuracy of our method was evaluated against visually scored recordings and obtained an average accuracy, F1 and Cohen Kappa score of 94.2%, 86.4%, and 83%, respectively, across all epochs. Our study contributes to the automation of EEG artifact detection by presenting a straightforward initial method for identifying contaminated epochs based on the spectral slope of a single EEG channel in rodent recordings.

## I. Introduction

Electroencephalography (EEG) is a non-invasive method which measures transmembrane currents resulting from the collective activity of cortical neurons detected by electrodes placed on the scalp [1]. Neural activity travels through tissue and bone before it is recorded, making the signal vulnerable to contamination as well as weakening the signal’s amplitude [1]. The challenge faced when analysing these signals is accurately separating measurements of neural activity from signals that are contaminated by external sources, whether these are physiological (ocular, muscular) or technical (hardware, body or cable movement). Components that are not directly produced by neural activity are known as ‘artifacts’. A gold-standard EEG processing pipeline to remove artifacts currently does not exist as it is difficult to define characteristics of artifacts that clearly distinguish these components from neural activity, and even more difficult to prove perfect signal extraction following data cleaning [2]. Therefore EEG data rejection is often extremely time-consuming and subjective, relying on multiple researchers visually assessing a recording and subsequently considering their collective assessment [3]. There are a large number of data cleaning pipelines available, however, it has been reported that several of these techniques can introduce artifacts themselves [4], and it can be difficult to know which technique to use first, or which method is suitable to apply to one’s data depending on the type of artifact present. For example, state-of-the-art techniques such as Independent Component Analysis (ICA) rely on multiple EEG channels to separate a multivariate signal into independent components [5], in addition to requiring in depth knowledge of EEG activity that is typically considered an artifact [6].

### A. Power Analysis of EEG signals

The power spectrum is a well-known linear analysis approach which transforms EEG signals from the time domain to the frequency domain through the Fourier Transform [7]. This transform decomposes the signal into its individual frequency components, which allows for quantification of the amount of activity in selected frequency bands [7]. In EEG analysis, distinct frequency bands have become associated with particular behavioural states, with the most commonly studied bands in animal studies including delta (1 - 4Hz), theta (8 - 12Hz), beta (12 - 30Hz) and low gamma (30 - 60 Hz) [8]. Particular frequencies have become associated with specific behavioural states, such as low-frequency delta activity during non-rapid eye movement (NREM) sleep, and high frequency activity during wakefulness (such as low gamma). These oscillations are easily detectable in power spectra plots as they have characteristic ‘peaks’ in the corresponding frequency bands [8]. Recent work has suggested that alongside these periodic peak components, there is an aperiodic ‘background’ component that is an important measurement of underlying neural activity [8]. This background activity can be quantified by the gradient of the power spectrum, known as the spectral slope, and has been shown to change dynamically with age [9], during sleep [10], under anesthesia [11] and in neurological disorders [12]. The aperiodic component of EEG signals have a characteristic 1*/f* power-law shape, exhibiting time- scale invariance, meaning that no particular temporal scale or frequency band dominates EEG activity. This distribution is present at both the cellular level (from neural membrane potentials) [13] and macroscopic EEG recordings, suggesting scale-free activity at both levels [14]. The 1*/f* component of the power spectrum may be described by the following power law function:

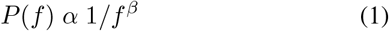

where *P* is power, *f* is frequency and *β* is the scaling exponent parameter describing the steepness of the aperiodic component’s slope. Equivalently, the equation can also be written as:

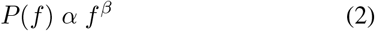

where the exponent for the decaying case is negative. When the power spectrum is plotted on a logarithmic scale this results in a linear function where the slope is the gradient of the linear function. As this function follows the power-law shape, the gradient of the spectral slope linear function is typically negative.

### B. Artifact Identification

Methods using the spectral slope to identify and remove noisy epochs using slope thresholds have been described, however, it has not been well-characterised as a stand-alone data cleaning method [2]. Furthermore, comparisons of spectral slopes between artifacts and clean epochs have not been analysed, making it unclear how to define the slope threshold.

In addition to this, studies have also reported variability in spectral slopes between individual recordings, suggesting there may be subject specific variations that should be considered before applying the same threshold to all recordings [15]. A small degree of variability may be expected due to how the recording device is attached or variations in the location of electrode placement across individuals, however this is not expected to affect the choice of spectral slope thresholds between individuals. As the power spectra can be used to identify dominant frequencies in a signal, it can also be used as a first pass preprocessing method to identify which epochs in a recording contain external artifacts. To assess whether the artifacts are unique to individual recordings or consistent across the population, we analyse the accuracy of different slope values within individual recordings.

## II. Methods

### A. Data Collection

EEG recordings were collected from 5 Long-Evans rats over a period of 24 hours, and implanted with 16-channel EEG surface grids with two integrated electromyogram (EMG) leads (Custom H16-Rat EEG16 Functional-NeuroNexus, United States). Electrode implants were connected to wireless amplifiers and recorded using a TaiNi wireless multichannel recording system (Tainitec, United Kingdom) [16] at a sampling rate of 250.4 Hz. EEG electrodes were placed over the following right and left cortical areas: primary somatosensory cortex trunk (S1 Tr), frontal association secondary motor cortex (M2 FrA), secondary anterior motor cortex (M2 ant), frontal anterior primary motor cortex (M1 ant), secondary visual mediomedial cortex (V2 ML), monocular primary visual cortex (V1 M) and primary somatosensory cortex hindlimb and forelimb region (S1HL S1FL). All spectral slope calculations are from the M2 anterior right channel (M2 ant right) as this channel was the most consistent across recordings visual inspection.

### B. Visual Artifact Identification

To compare how accurately the spectral slope algorithm is able to identify these artifacts, recordings were visually scored to label epochs with artifacts present (such as the signal disruption in Fig. 1B) and those without. Artifact identification was performed for all 24 hour recordings by the same individual, and epochs were labelled as contaminated if there were any technical or physiological artifacts present that would prevent the sleep stage classification (rapid eye-movement (REM), non-rapid eye movement (NREM) or wakefulness) of that epoch from being determined. Epochs in four out of five recordings were also manually classified as rapid eyemovement (REM), NREM and wakefulness to determine if the presence of artifacts is most prevalent during one particular sleep stage. The defining criteria for each stage are defined by the American Academy of Sleep Medicine (AASM) [17].

**Fig. 1:**
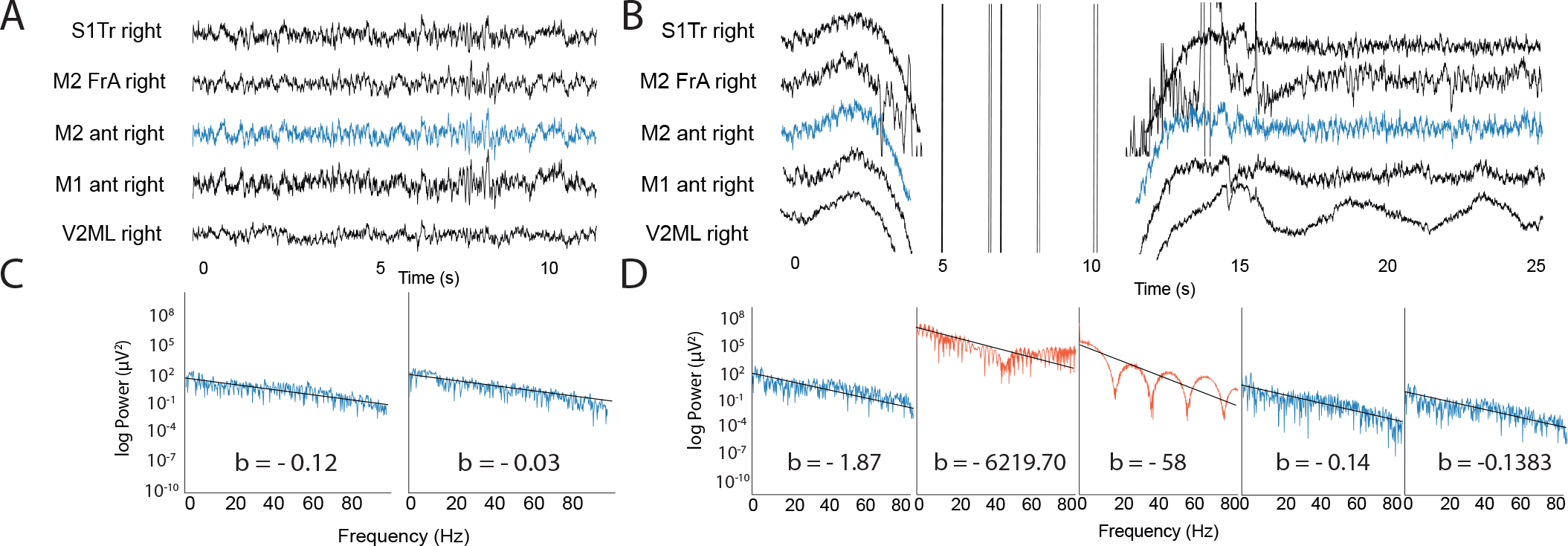
(A) Raw EEG traces from 2 consecutive 5 second time bins with no artifacts present (B) and with artifacts present between 5 and 15 seconds. Note: the channel selected for spectral analysis is shown in blue. (C, D) Power spectra (blue) and spectral slopes (black) are shown for 5 s epochs with no artifacts present (blue) and with artifacts present (orange) for traces in A and B.

### C. Data Preparation

A bandpass butterworth filter (0.2 - 100Hz) was applied to the full-length raw data. The recording was subsequently separated into 5 second epochs (with each recording having a total of 17280 epochs). The time bin length of 5 seconds is commonly used in EEG analysis and should prevent long periods of data being discarded due to short artifacts being present whilst also ensuring appropriate sampling length for windowing [18]. Prior to spectral slope calculations, epochs with voltage values that exceeded 3000mV were removed as this is beyond the range of physiological activity.

### D. Spectral Slope Calculations

Epochs were Hanning-tapered, and Fourier transformed using the SciPy welch method (with 50% overlapping windows) to obtain the spectral density at each frequency. A first linear least-squares line was fit to the logarithmic power spectra of each epoch using the numpy polyfit function (with a degree of 1) ranging between 1 and 48 Hz (to avoid any power line artifacts at 50Hz). This method aims to find slope values that minimise the sum of the differences between the observed EEG power values and the least-squares line.

### E. Model Evaluation

To evaluate how accurate the spectral slope is at differentiating between epochs with and without artifacts, we used several metrics - the F1 score in particular - to account for the class imbalances between the number of clean and noisy epochs (with the average split across all recordings being 80% clean epochs and 20% noisy epochs). We calculated the accuracy, Cohen’s kappa, precision, recall and F1 scores of the spectral slope algorithm for different slope values and compared these measurements for each recording. Cohen’s kappa (*K*) scores measure the degree of agreement between two scorers and are primarily used for categorical data such as sleep scores [18]. K scores *x* ≤ 0 signify no agreement, 0.01 ≤ *x* ≤ 0.20 fair agreement, 0.41≤ *x* ≤0.60 moderate agreement, 0.61≤ *x* ≤0.80 substantial agreement, and 0.81≤*x* ≤1.00 perfect agreement [18].

## III. RESULTS

### A. Technical artifacts inflate the overall average power

To determine the effect artifacts have on the overall signal average, the average power spectra across all epochs for the selected channel was calculated before spectral slope preprocessing (Fig 2A) and compared to the average power spectra average after spectral slope values less than -8 *μV* ^2^*/Hz* were removed (Fig 2B). The influence of these artifacts on the overall power values is striking, as reflected in the significant increase in mean power to 10^6^ *μV* ^2^ before artifact removal, compared to 10^2^ *μV* ^2^ after artifact removal (Fig 2A, B). Along with this, prior to preprocessing the data follows similar spectral slope trends, but after artifact removal there is an emergence of distinct individual spectral patterns (such as periodic component ‘bumps’ appearing in recording 2 and 3) (Fig 2B). It is therefore evident that without the removal of noisy epochs, the objective of understanding patterns of neural activity with spectral analysis is impractical as the presence of these artifacts conceals any periodic or aperiodic spectral pattern that may be present.

**Fig. 2:**
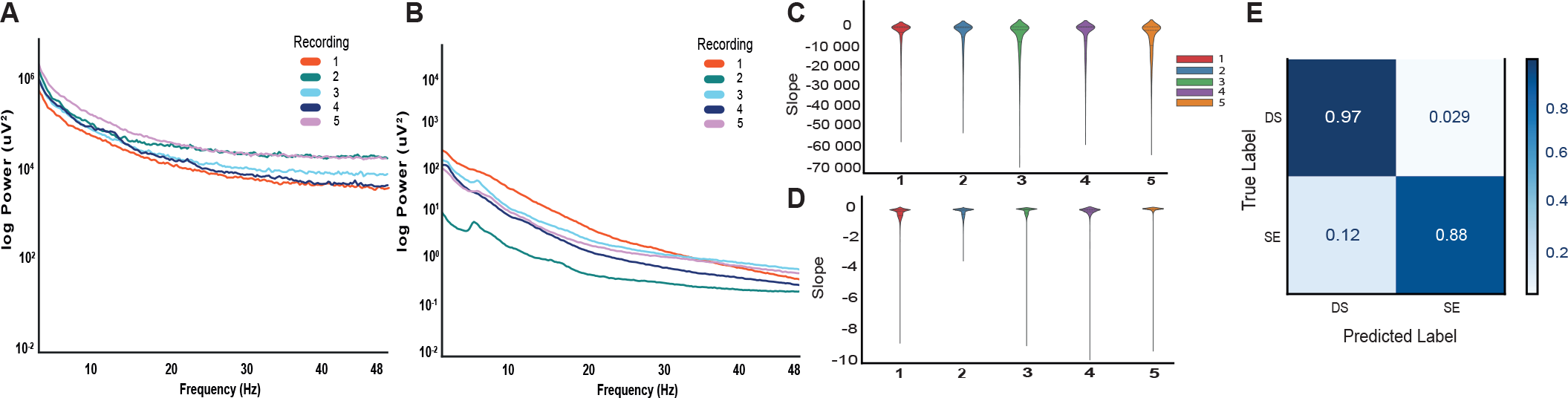
Power spectra averages across all epochs without any spectral slope preprocessing (A) and after excluding epochs with a spectral slope less than -8 (B). (C) Violin plots comparing the distribution of spectral slopes of visually scored epochs labelled as ‘noisy’ in five separate recordings, compared to those labelled as clean (D). Lower quartile, median and upper quartile values are shown by dotted lines. The range of slope values is particularly evident between noisy and clean epochs. (E) A confusion matrix showing the predictions made by the spectral slope for one recording.

### B. Testing individual variance

We evaluated how effective various spectral slope values were at classifying an epoch as artifact-free or contaminated for each recording to determine the most appropriate spectral slope threshold (Fig 2C - E). To identify appropriate slope values to compare, we analysed the distribution of values for epochs labelled as clean and noisy, with a particular focus on overlapping values between the two classes. For clean epochs, we observed median spectral slope values of -0.671, -0.118, -0.432, -0.082, and -0.137 for recordings 1, 2, 3, 4, and 5, respectively, along with corresponding mean values of -1.65, -1.04, -1.80, -0.54, and -1.28 (see Fig 2C). In contrast, the median values for the noisy epochs were considerably different, measuring -285.28, -403.42, -1440.06, -175.75 and -1741.68 whilst the mean values were -2911.18, -4063.28, - 5424.08, -4897.47 and -6071.80 for each respective recording (Fig 2D). We tested spectral slope thresholds of -6, -8, - 10 and -12 based on the majority of clean epochs having clean epochs greater than -4, and found that a threshold of -8 obtained the highest accuracy scores, achieving scores of 95%, 0.87%, 0.96%, 0.98% and 0.95% for recordings 1, 2, 3, 4 and 5, respectively. The F1 scores were 93%, 0.85%, 0.89%, 0.78% and 0.87% whilst K scores were 92%, 0.83%, 0.71%, 0.87%, and 0.84% for recording 1, 2, 3, 4 and 5, respectively, which fall into the category of perfect agreement. Recording 3 obtained only a substantial agreement score as it was one of the recordings that had spectral slopes with values below -8 for epochs labelled as clean (Fig D). This could also be the result of noisy epochs being labelled as clean during manual scoring, illustrating the issue of human bias during manual scoring. Overall, the combination of these evaluation methods suggest that this method can effectively identify epochs containing artifacts, however, the effectiveness of this method is dependent on an appropriate spectral slope threshold.

### C. Investigating Artifact Sources

Finally, to determine the possible cause of the artifacts identified by the spectral slope, we quantified the sleep scores in the epochs adjacent (before and after) to the epoch labelled as noisy (Table 1 & 2). If the artifacts are primarily present during one particular sleep stage, this would indicate that they could be the result of behaviour that only occurs in that sleep stage (such as chewing or movement during wakefulness) [20]. Epochs were classified as noisy most commonly during wakefulness, with 73.13% of noisy epochs having adjacent epochs classified as wakefulness in comparison for 23.51% and 3.35% for NREM and REM, respectively (Table 1). This is compared to the percentage splits for clean epochs per sleep stage being 48.06%, 47.14% and 4.96% for wakefulness, NREM and REM. The increased frequency of noisy epochs during wakefulness suggests that they could in fact be the result of behaviour that is more common to this state. However, it does not completely explain the source of artifacts as there are still quite a few ‘noisy’ epochs in both REM and NREM, and REM in particular is characterised by an absence of body movement [19], and as a result one would expect very few or no REM epochs contaminated with artifacts. Whilst there are very few REM epochs contaminated with artifacts (3.35% of noisy epochs, Table 1), this is still consistent with the percentage of clean epochs (4.96%, Table 2).

**TABLE 1:** Number of Contaminated Epochs by Sleep Stage.

| Recording | Wake | NREM | REM | Total |
| --- | --- | --- | --- | --- |
| 1 | 1632 | 984 | 78 | 2694 |
| 2 | 1669 | 513 | 132 | 2314 |
| 3 | 1585 | 268 | 71 | 1924 |
| 4 | 1682 | 343 | 21 | 2046 |
| **Total** | 6568 | 2108 | 302 | 8978 |
| **Percentage** | 73.15% | 23.51% | 3.35% | 100% |

**TABLE 2:** Number of Clean Epochs by Sleep Stage.

| Recording | Wake | NREM | REM | Total |
| --- | --- | --- | --- | --- |
| 1 | 6898 | 6390 | 324 | 13612 |
| 2 | 6849 | 6527 | 956 | 14332 |
| 3 | 5970 | 7254 | 1197 | 14421 |
| 4 | 7013 | 5924 | 284 | 13221 |
| **Total** | 26730 | 26195 | 2761 | 55586 |
| **Percentage** | 48.06% | 47.14% | 4.96% | 100% |

## IV. Discussion

EEG signals are widely used in clinical and research settings to better understand physiological processes underlying cognition and development [20]. EEG electrodes are however susceptible to unwanted noise from sources external to neural activity, and the manual identification of this noise is incredibly time consuming. We have proposed a first pass artifact detection method to identify epochs containing artifacts from EEG recordings using the spectral slope, requiring only one EEG channel to identify these epochs, which obtains an overall accuracy, F1 and Cohen Kappa scores of 94.2%, 86.4% and 83%. This method could, however, also be extended to include all electrode channels to similarly identify whether specific channels are more affected by artifacts or to identify the most suitable channel for further analysis, such as automated sleep scoring.

### A. Limitations

Precaution should be taken when selecting the epoch length as this will determine how much of the recording is removed. If the chosen epoch length is too long, this could result in large portions of the data being removed even if an artifact is only present for a very small portion of that time bin. However, if the epoch length is too small this may in turn affect the windowing function in the power spectra calculations. Furthermore, it is important to note that the origin of these artifacts is unclear, and the spectral slope is unable to identify artifacts such as eye-blinks as these share the same spectral slope range as typical activity [21]. Consequently, distinguishing these artifacts using a spectral slope threshold is challenging. As a result, our suggested spectral cleaning method does not negate the need for preprocessing methods such as ICA to identify artifacts in the physiological range. It can rather be applied prior to ICA to identify epochs that should be removed, and save time by preventing visual scoring of these epochs.

### B. Future Work

We analysed recordings obtained using wireless, as opposed to tethered, technology in rats. Wireless recording devices are generally considered superior due to the risks of injury and artifacts resulting from the attached cables from tethered devices being absent [22]. However, the spectral slope method can also be applied to recordings from tethered animals, and is also effective at removing similar artifacts [23]. The use of one channel to identify artifacts means this technique is not dependent on the electrode set up. Whilst the spectral slope is scale and time invariant, the slope is reported to be steeper (more negative) in electrocorticography (ECoG) recordings in comparison to local field potentials (LFP) [24]. It would therefore be interesting to perform the same analysis across multiple recording techniques (LFP and ECoG) and species to compare whether the spectral slope threshold is dependent on the recording method or type of species. The advantage of the spectral slope method is that it does not consider differences between defined frequency bands that are typical in spectral analyses, but rather the change in power over a large range of frequencies [8]. As a result, it could be easily translated to human studies where relevant individual frequency bands are slightly different to those analysed in rodent studies but exist in the same overall frequency range (1 - 48*Hz*) [25].

## Conclusion

Our study advances the automation of EEG artifact detection by providing a straightforward initial approach for identifying contaminated epochs using the spectral slope of a single EEG channel. This approach has the potential to significantly reduce the time required for visual identification of artifacts and carries no risk of introducing additional artifacts. However, it is essential to exercise caution when choosing the epoch length, taking into account factors such as the sampling rate and the percentage of overlapping windows. Choosing an appropriate slope threshold is the determinant of the algorithm’s classification accuracy. We found that a fixed spectral slope threshold of -8 *μV* ^2^*/Hz* was successful in identifying noisy epochs across all EEG recordings. To expand the applicability of this method, further analysis should be repeated in LFP and ECoG animal recordings in addition to human EEG studies to determine whether the same threshold can be applied to diverse recording methods.

## Acknowledgment

*Alfredo Gonzalez-Sulser and *Javier Escudero contributed equally to this work.

## Notes

### Competing Interest Statement

The authors have declared no competing interest.

